# Supervised Rank aggregation (SRA): A novel rank aggregation approach for ensemble-based feature selection

**DOI:** 10.1101/2022.02.21.481356

**Authors:** Rahi Jain, Wei Xu

## Abstract

**Background:** Feature selection (FS) is critical for high dimensional data analysis. Ensemble based feature selection (EFS) is a commonly used approach to develop FS techniques. Rank aggregation (RA) is an essential step of EFS where results from multiple models are pooled to estimate feature importance. However, the literature primarily relies on rule-based methods to perform this step which may not always provide an optimal feature set.

**Method and Results:** This study proposes a novel Supervised Rank Aggregation (SRA) approach to allow RA step to dynamically learn and adapt the model aggregation rules to obtain feature importance. The approach creates a performance matrix containing feature and model performance value from all models and prepares a supervised learning model to get the feature importance. Then, unsupervised learning is performed to select the features using their importance. We evaluate the performance of the algorithm using simulation studies and implement it into real research studies, and compare its performance with various existing RA methods. The proposed SRA method provides better or at par performance in terms of feature selection and predictive performance of the model compared to existing methods.

**Conclusion:** SRA method provides an alternative to the existing approaches of RA for EFS. While the current study is limited to the continuous cross-sectional outcome, other endpoints such as longitudinal, categorical, and time-to-event medical data could also be used.

## Introduction

A high dimensional data has challenges associated with model fitting, generalizability [1], and computation complexity [2,3], which prevents modeling by many classic statistical techniques. Feature selection is an important component in high dimensional data analysis domains like genomics [4] and radiomics [5], as it helps reduce the dimensions of the dataset. Literature provides many techniques to perform feature selection. However, these techniques could be categorized based on their feature selection (FS) approach (Figure 1). One broad category of FS techniques uses only expert or domain knowledge to perform feature selection [6,7]. These techniques work in scenarios with few features without interaction among features and are well known in the research domain [8]. Another broad category of FS techniques combines expert or domain knowledge with data [9,10]. FS techniques designed in the Bayesian framework incorporate prior knowledge in the feature selection process [9].

**Figure 1:**
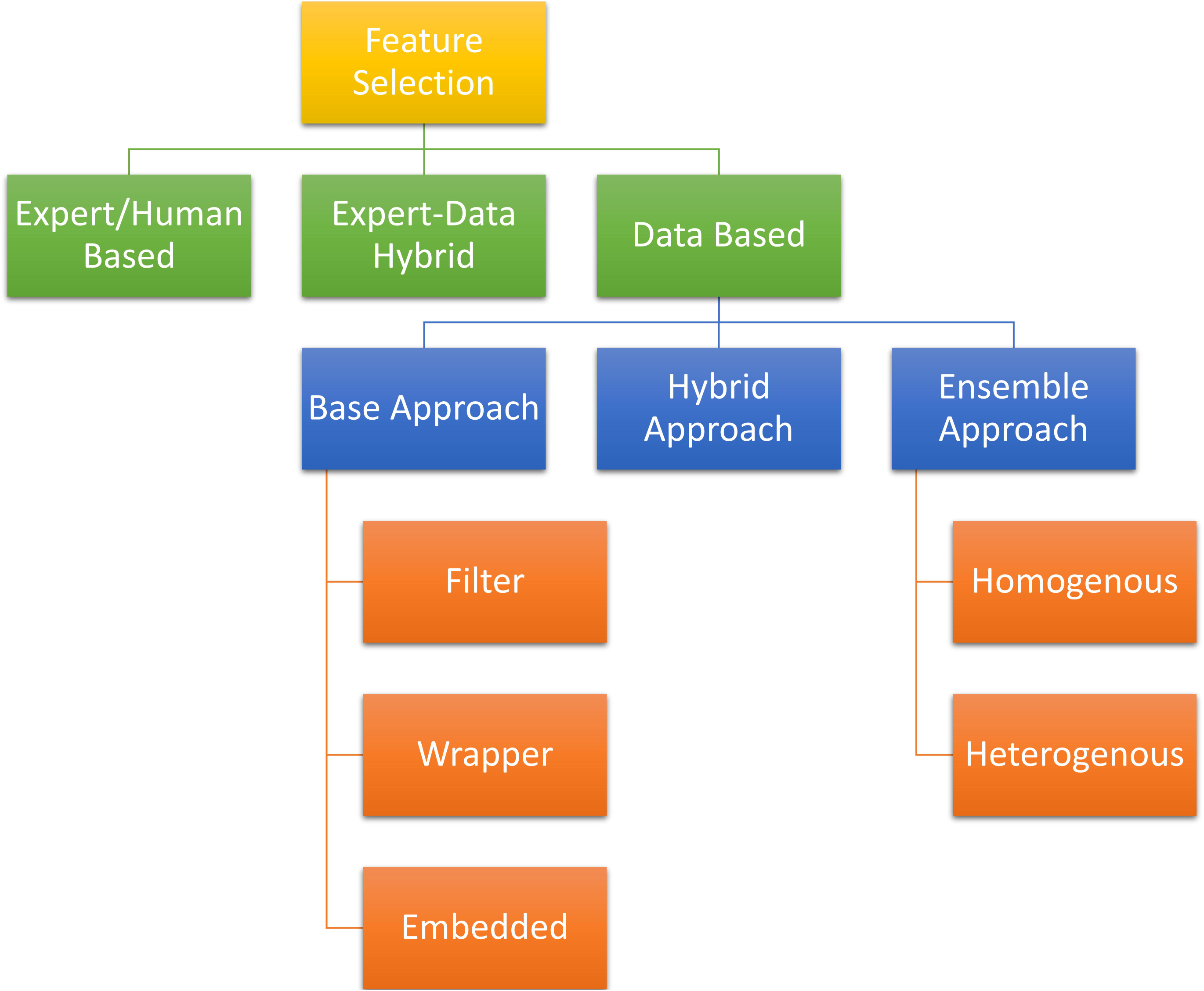
Different feature selection approaches.

The third and major category of FS techniques relies on the dataset to perform feature selection and is referred to as data-based FS techniques in this paper. These techniques are sub-categorized into Filter, Wrapper, and Embedded FS techniques [11,12]. Filter methods select features based on internal data structures like association [13] and information gain [14,15]. Wrapper methods evaluate multiple subsets of features iteratively by building models to get the feature subset, which achieves the best performance [16–18]. Embedded methods build the model that simultaneously performs features selection [19–22].

Literature suggests different approaches to use the FS techniques for FS. These approaches can be categorized into base, hybrid, and ensemble approaches. In the base approach, a single FS technique is used. In the hybrid approach, multiple FS techniques are used in a sequence to perform feature selection [10,23]. Commonly, a filter based FS technique is used as coarse FS followed by a wrapper or an embedded based FS technique for final FS [23]. Some approaches create a sequence by combining expert based FS with other FS techniques [10].

In an ensemble approach, instead of a single model, multiple models are created from the same dataset. The performance of features from these models is pooled and ranked based on their relevance. Finally, the relevant features are selected based on the cut-off of importance. Two approaches can be used to generate multiple models, namely homogenous ensemble approach and heterogeneous ensemble approach [24–26]. In a homogenous ensemble approach, multiple datasets are created from the same data by sub-setting the samples, features, or both followed by using a single technique to build the model on each of these datasets [25]. In a heterogeneous ensemble approach, a single dataset is modeled using different techniques to generate multiple models [26]. An ensemble approach could perform better than single model approaches [27].

In an ensemble approach, one of the essential steps is to pool together the performance of features obtained from different models and is referred to as rank aggregation (RA) in this study. The performance metric used for RA varies across the studies, like model estimates [8,21] and goodness of fit [8]. Literature provides various techniques to aggregate the feature performance obtained from different models, but these techniques mainly rely upon a pre-defined rule to aggregate the performance of features, i.e., rule-based rank aggregation approaches. Commonly used methods to aggregate the performance of features is to find the mean, median or Robust rank aggregation (RRA) performance of the feature across all models [28] [29]. However, they cannot learn from the data about the RA rule dynamically and may even be sensitive towards extreme values like mean values.

In high-dimensional data analysis, the performances of rule-based analysis have been challenged by machine learning (ML) based approaches like supervised learning. ML-based approaches are considered effective in the dynamic and complex environment as compared to rule-based approaches because ML creates dynamic rules by learning and adapting to the existing environment [30]. In the case of ensemble FS, the data structure is dynamic and varies across datasets, so it may not always be possible for a predefined rule to give optimal results for all the scenarios [30,31]. Thus, it is desirable to explore the application of ML in all steps of ensemble FS owing to its dynamic learning characteristics. ML approaches like supervised learning are well established in the model building step [32], but no supervised learning approach is designed for the RA step.

This study proposes a novel perspective to perform RA using the supervised learning approach of the ML called supervised rank aggregation (SRA). First, SRA creates a performance matrix that contains the performance of all features in all the models as the input and the performance of each model in achieving the final data analysis goal as the label. Then, supervised learning is used to find the relative rank or performance of features based on their potential to help achieve the best performance in the final data analysis.

SRA based ensemble feature selection (EFS) is highly innovative in many ways. Firstly, perspective is unique as it pools and ranks features dynamically rather than using fixed rules for EFS. Secondly, it provides a unique application of supervised learning models as they replace the static rule-based RA approach with a dynamic rule-based RA approach. Thirdly, it is versatile, which allows its integration with existing ensemble methods.

This paper provides the “Methodology” section to explain the SRA based EFS. Then, its performance is compared against existing rank aggregation methods used in EFS for simulations and real studies in the “Simulation Studies” and “Real Studies” sections. Finally, we summarize and provide future directions for research in the “Conclusion and Discussion” section.

## Methodology

SRA methodology is developed to integrate the supervised learning in the rank aggregation step of the ensemble learning (Figure 2). A dataset of sample size, *n*, with given input feature space, *p*, and an outcome is fed into the EFS process, where multiple models are created either by creating multiple bootstrapped datasets from the original dataset (homogenous approach) or by using multiple modeling techniques (heterogeneous approach). Then a performance matrix is created from these multiple models by extracting feature performance and model performance. A supervised learning algorithm is trained on this performance matrix, and feature importance obtained from the algorithm is used as the final feature ranking or importance. Finally, the features are selected based on an importance cut-off obtained from a predefined threshold or an unsupervised ML algorithm. The proposed methodology is discussed below in more detail.

**Figure 2:**
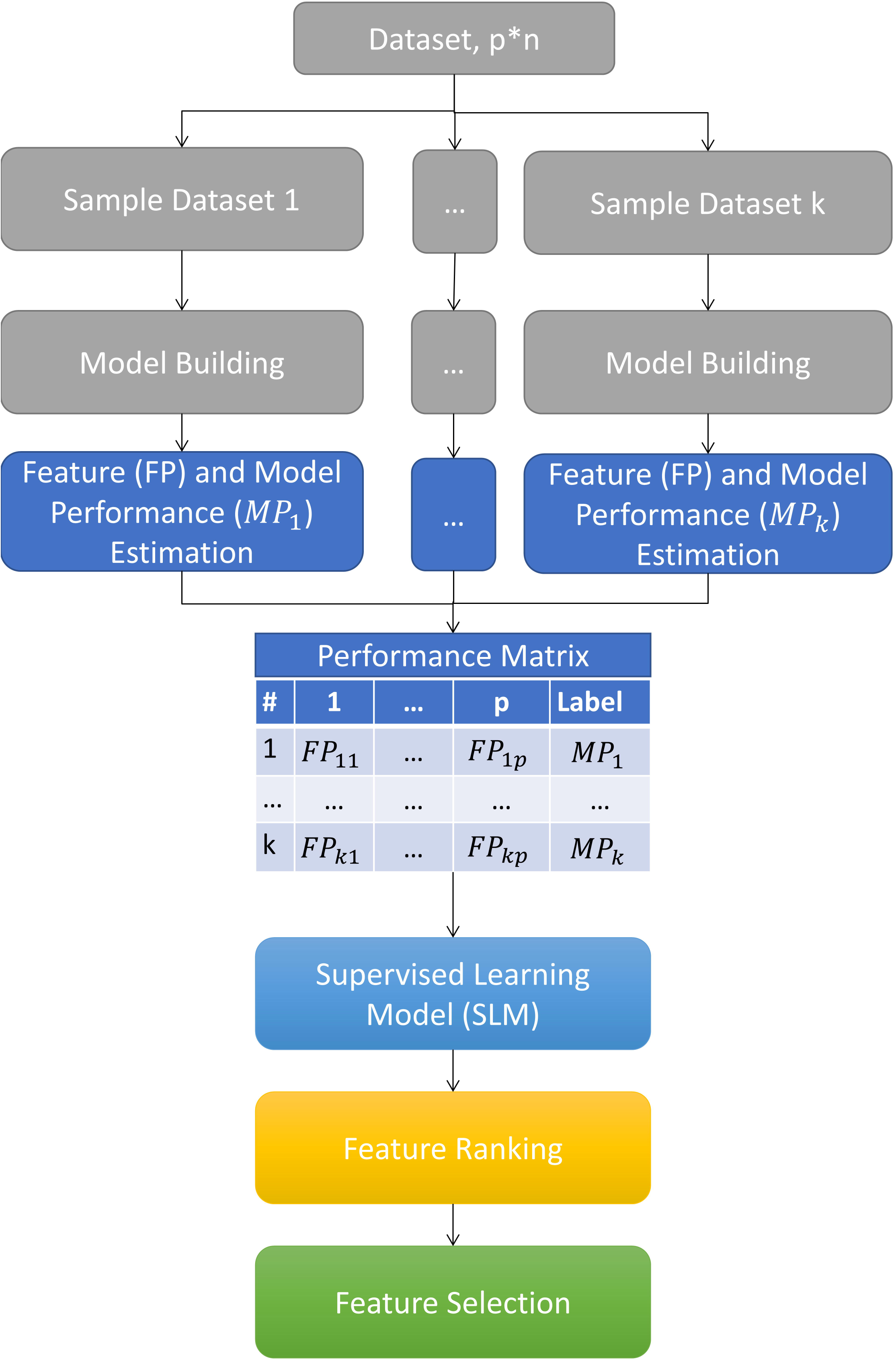
Graphical representation of SRA methodology based Ensemble feature selection.

### Generate multiple models

From the original dataset *D* of feature space *p*, outcome *y*, and sample size *n,k* randomly sampled datasets are generated by randomly sampling features without repeats *q*_*i*_ | *i* ∈ {1, …, *k*},1 < *q* ≤ *p*. All *k* sample datasets have a sample size of *n* by sampling with replacement from dataset *D*. A model *m* is created for every *k* sample dataset using any modeling technique.

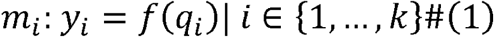

where, modeling technique used to prepare the *i*^*th*^ dataset model *m*_*i*_ will determine the function *f*. In this study, RIDGE regression is used as the modeling technique for building the models. Optimal hyperparameter values for each model are obtained from 10-fold cross-validation.

### Create Performance Matrix

A performance matrix *C* is prepared from *m* models containing *k* rows and *p* + 1 columns. The matrix contains feature performance, *FP* as the input features and model performance for study objective, *MP* as the outcome or label for all *m* models.

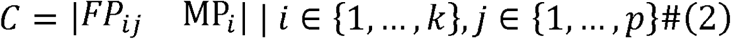

In the current study, model estimates are used as *FP* metric and predictive performance of a model on the left out bootstrap samples as *MP*. Accordingly, *MP* metric used in the study is inverse of root mean square error (RMSE).

### Supervised Rank Aggregation

A supervised learning model (SLM) is created from the performance matrix with *FP* of *p* features as predictors and *MP* as the outcome.

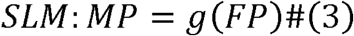

where, machine learning technique used for SLM will determine the function *g*. Currently, only ML techniques like penalized regression and decision trees which could provide feature importance, *fimp* in achieving the model performance could be used.

### Feature Selection

The importance for each feature is used to select target features *q*_*best*_. It is assumed that the features with more importance should be target features as they are more relevant in achieving higher model performance. In literature, the cut-off value for features is obtained by using a predefined threshold [8,21], rule-based threshold estimation [33], or unsupervised learning based threshold estimation [21]. A predefined threshold may require the tuning step to arrive at an appropriate cut-off value, which will give optimal results for a given scenario [8,21]. Rule-based methods may not always provide optimal results [30,31]. Thus, in this study, the K-means based unsupervised learning technique is used for obtaining the threshold cut-off as it will eliminate the need for tuning and dynamically adapt to the given scenario. Since clustering will be happening on a single dimension, hence high dimension limitation of K-mean clustering is avoided. K-means is used to cluster the features into two groups, and the features in the cluster with a higher mean *fimp* value are selected as final features *q*_*best*_. Pseudo Algorithm summarizes the complete SRA based ensemble feature selection algorithm.

#### Pseudo Algorithm: SRA based ensemble feature selection

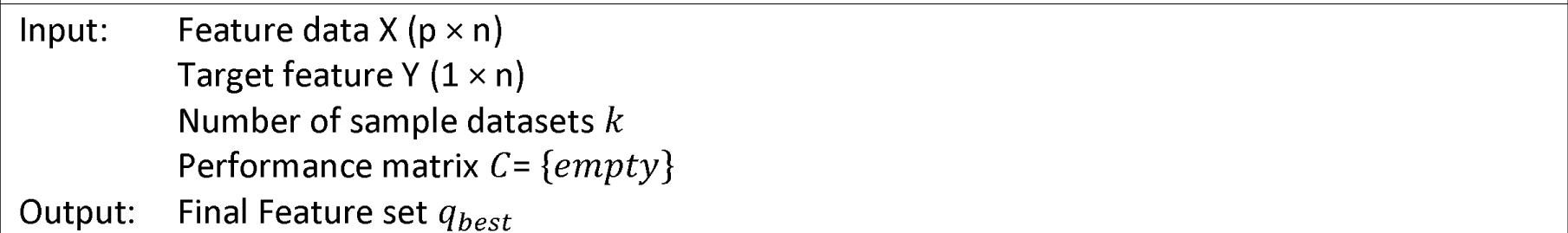
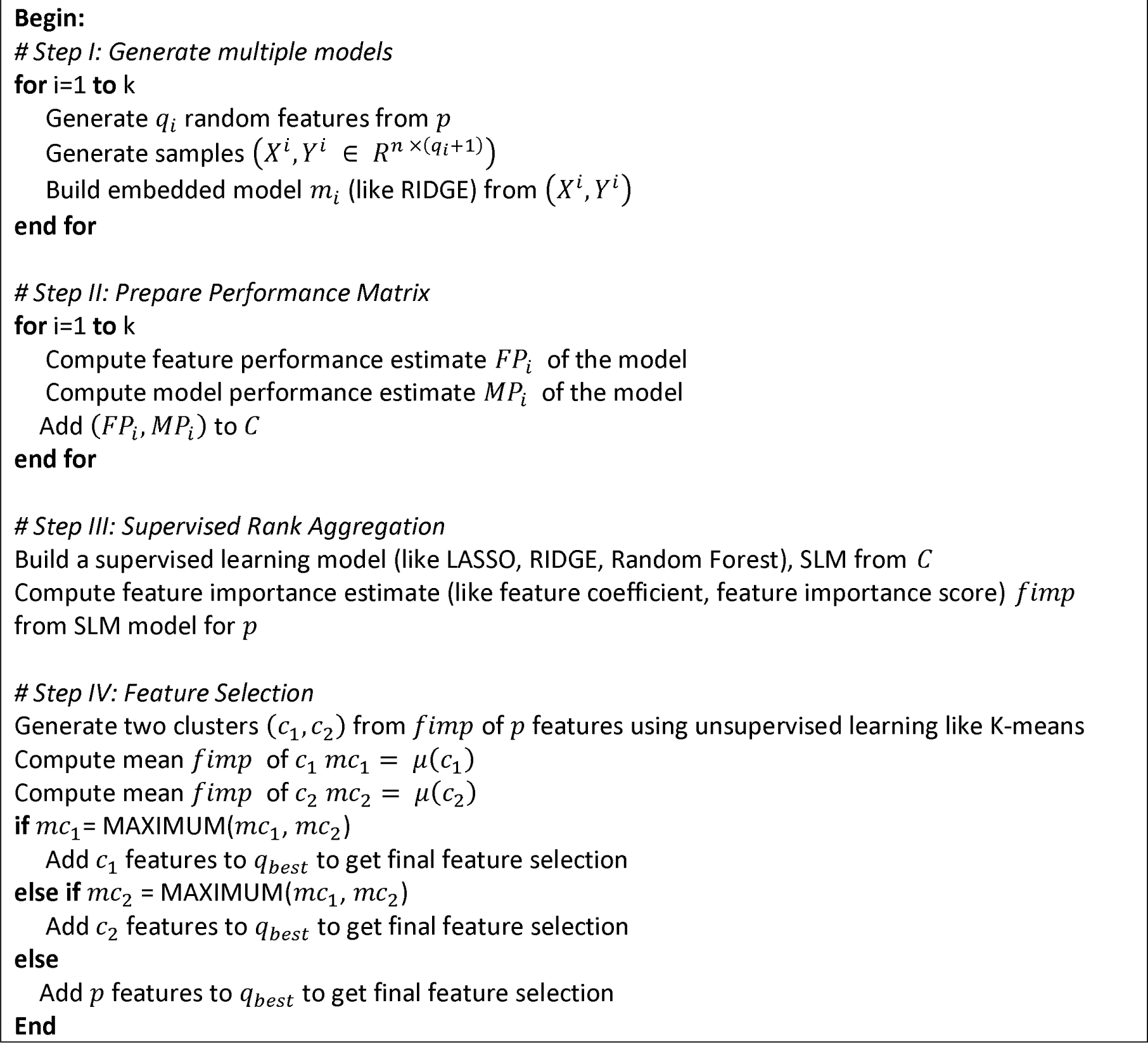

### Simulation Studies

We perform simulation studies to evaluate the proposed RA method and compare its performance with multiple other RA methods for EFS. The study generates high-dimensional feature space for marginal models using multivariate normal distributions. The study uses regression model 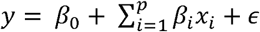 to provide a continuous outcome variable of simulated data with sample size, *n* for marginal models. *β* represents the effect of different features and intercept term on the outcome, *ε* ∼ (0, *σ*^2^) is the normally distributed error term and *x*_*i*_ ∼ *N* (0, 1) are normally distributed input features, *p*. Multi-collinearity is added between features using the covariance matrix as given below:

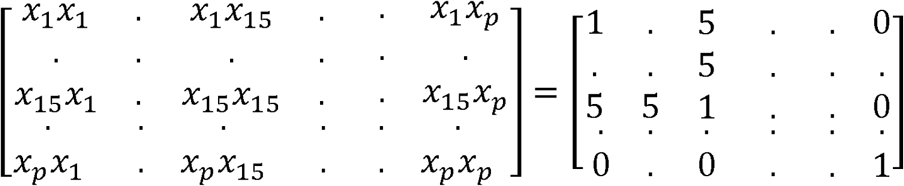

Multiple scenarios are simulated by changing *p, n, β*, the number of target features, and *k* (Table 1). Only true features are assigned a non-zero *β* value. We prepare homogenous ensemble models for feature selection. The dataset for each model is generated by randomly sampling two to *p* features from *p* feature space and sub-setting *n* samples from the original dataset with replacement. RIDGE is used to build models for each dataset. A penalized effect size of each feature obtained from *p*the RIDGE models is scaled using the absolute maximum value, which is used as a feature performance metric.

**Table 1:**
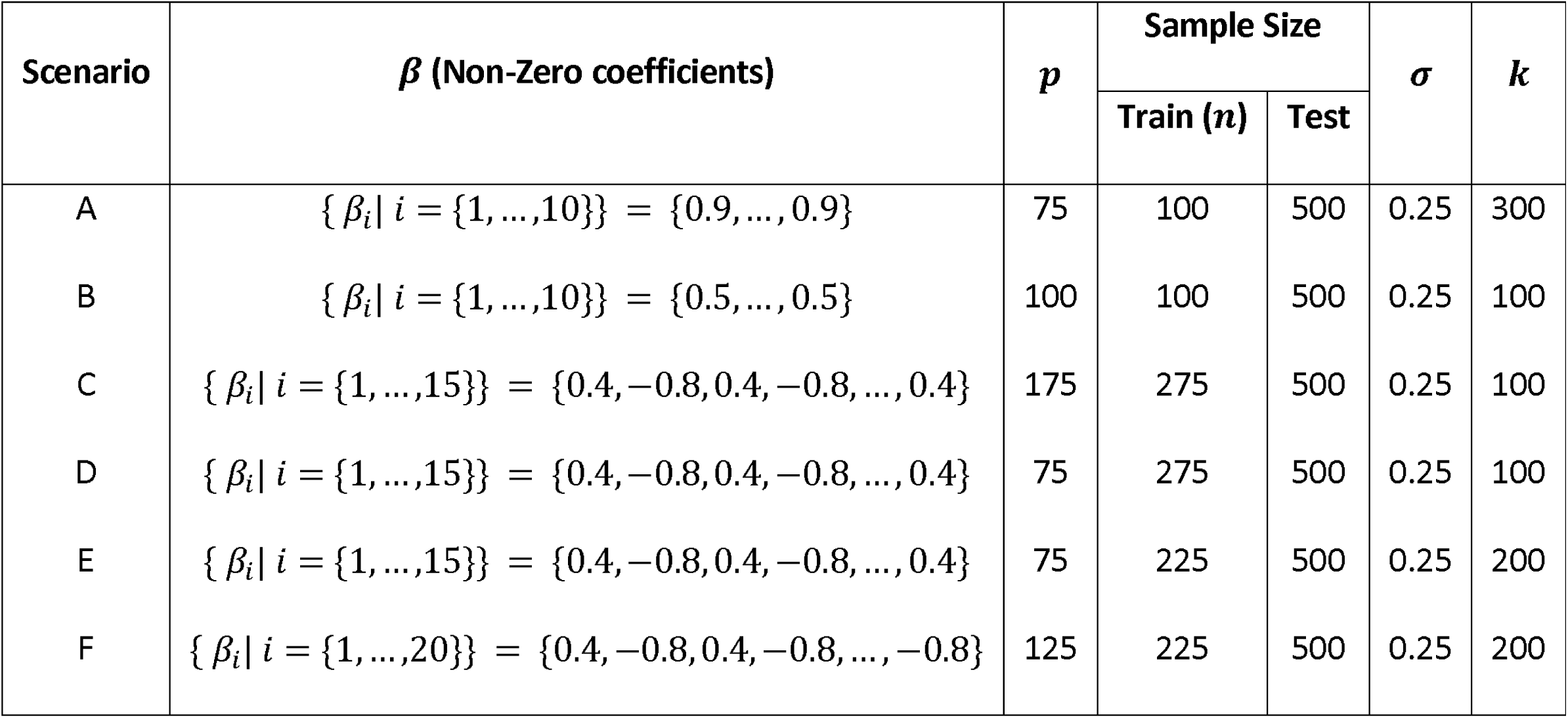
Description of the scenarios of simulation studies

Implementation of SRA is shown using three different supervised learning algorithms, namely LASSO (SRA-Lasso), RIDGE (SRA-Ridge), random forest (SRA-RF). While LASSO and RIDGE perform supervised learning using a linear combination of features, random forest performs supervised learning using a non-linear combination of features. A supervised learning model in each SRA is prepared using optimized hyperparameter values.

SRA performance is compared with existing rule-based RA methods, namely, mean based RA (MeRA), maximum based RA (MaRA), minimum based RA (MiRA), median based RA (MedRA), coefficient of variation based RA (CVRA), standard deviation based RA (SDRA), robust rank aggregation (RRA), t-test based RA (tRA), and Wilcoxon signed-rank test based RA (WRA). R 4.0.3 is used for the analysis. The study has used some inbuilt packages in statistical language R for the analysis like *glmnet* package [34] for LASSO and RIDGE, *randomForest* package [35] for random forest, and *RobustRankAggreg* package [29] for RRA.

The different RA methods are evaluated for their ability to select target features, discriminate between target and noise features, and predictive performance of the models built using selected features. We use the F1 score for feature discrimination ability evaluation and inverse RMSE for the test data for the predictive performance evaluation. RIDGE is used to build the final model from the selected features for predictive performance evaluation. Ten trials are performed for each scenario.

Table 2 results suggest that all methods can select some target features under all scenarios, but SRA-Ridge consistently outperformed rule-based RA methods. SRA-Ridge selected almost all the target features in all scenarios. The performance of the other two SRA methods is at par with existing RA methods. Further, the results suggest that SRA-Ridge has a better or at par feature discriminative ability than other methods. Thus, SRA-Ridge not only selects target features but is also good in rejecting noise features as compared to other methods. The results suggest that SRA could be a good candidate to select target features.

**Table 2:**
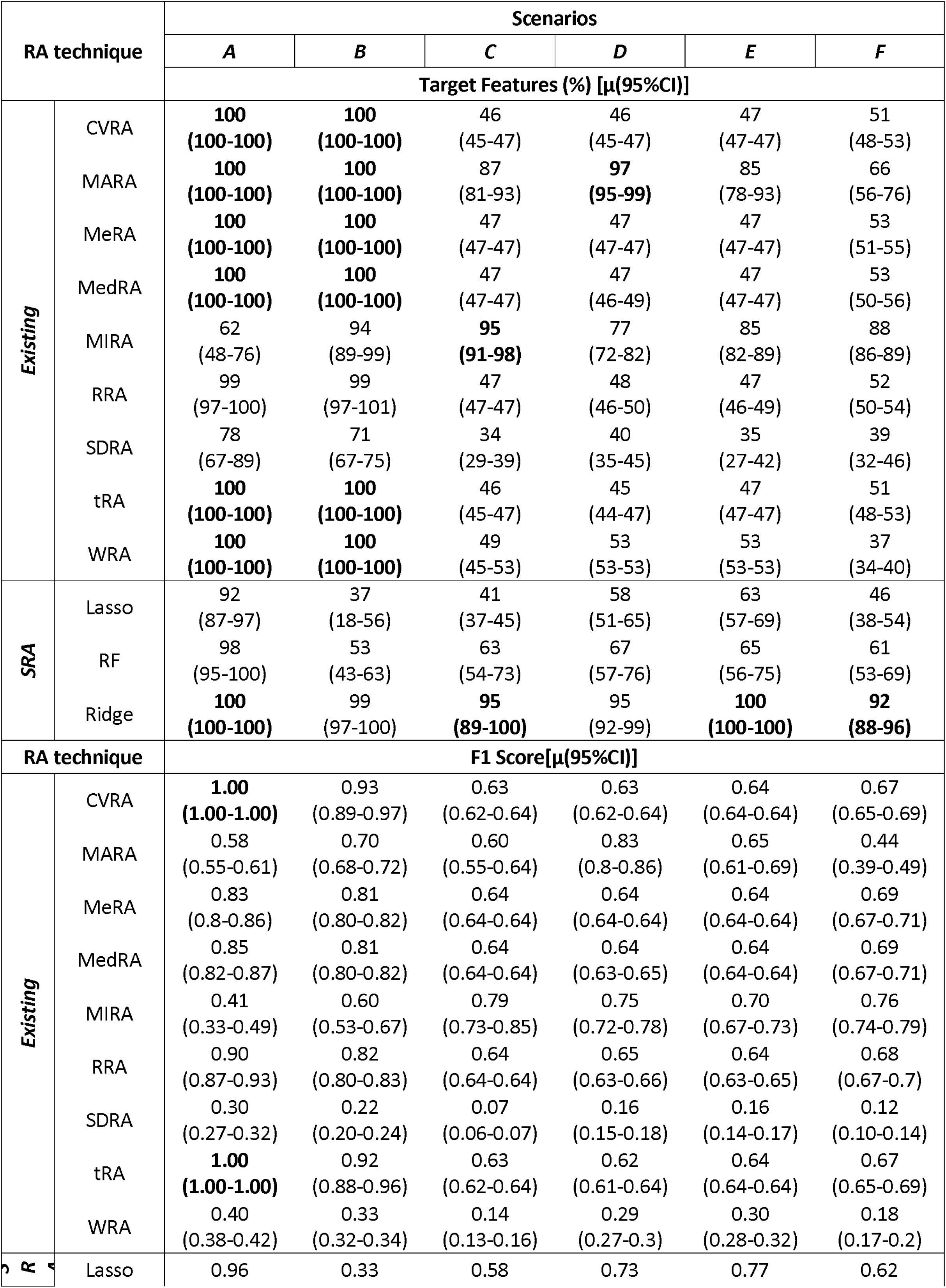
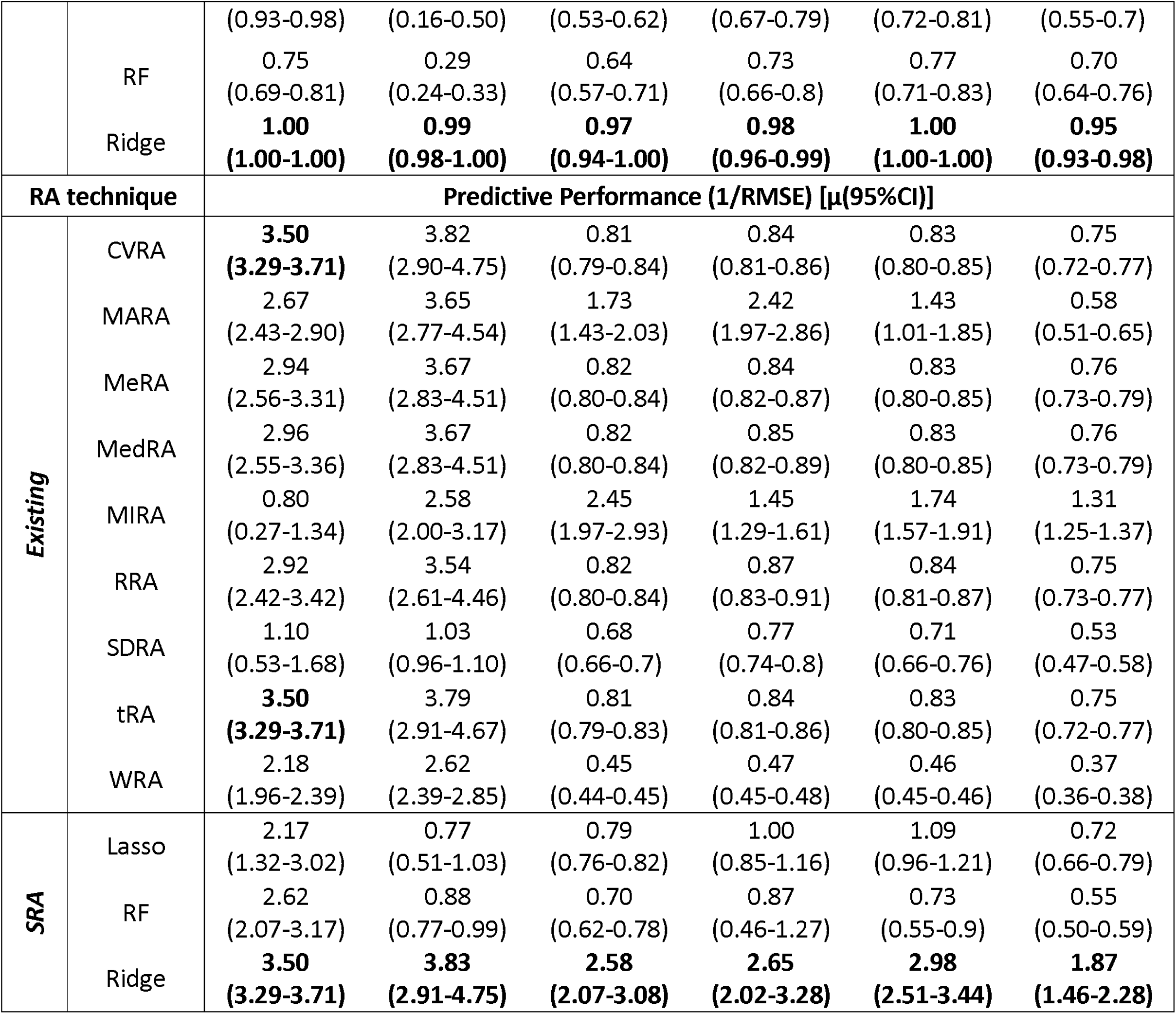
Comparison of model performance between SRA methods and Existing methods under six scenarios in terms of target feature selection, feature discrimination ability (F1 Score) and outcome prediction (1/RMSE)

Further, SRA-Ridge based selected features can build good predictive models and consistently outperformed rule-based RA methods (Table 2). These findings suggest that the SRA may provide better or at par prediction performance than existing methods. Further, SRA could enhance the performance of ensemble-based approaches in high-dimensional settings.

### Real Studies

Three real studies are analyzed to compare the performance of SRA and existing RA methods. Study I is Community Health Status Indicators (CHSI) study that collected US county data (n=3141) containing 578 features to understand non-communicable diseases [36]. Study II is National Social Life, Health and Aging Project (NSHAP) study that collected aged Americans data (n=4377) containing 1470 features to understand their health and well-being [37]. Study III is the DNA methylation data (n=27578) containing 108 samples to understand its relationship with human age [38,39].

Table 3 shows the final cleaned dataset for these three studies used for analysis. Features and samples are filtered to remove highly correlated features, non-continuous features, missing values, and very low standard deviation. The final cleaned dataset is randomly split into training and test dataset. The test dataset is used to evaluate the predictive performance of the features selected by different RA methods. The study uses inverse RMSE as the predictive performance metric. The mean performance of ten trials is used for comparison between RA methods. In the cases of Study I and Study II, 100 ensemble models are created, while in Study III, 1000 ensemble models are created.

**Table 3:**
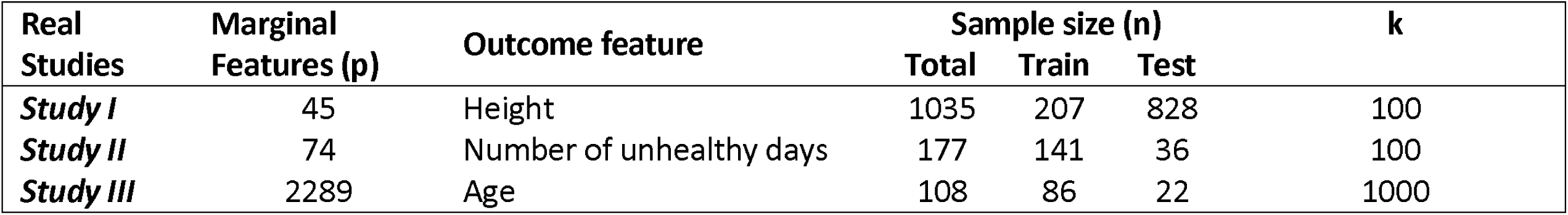
Summary of the real datasets

The results from Table 4 suggest that SRA methods provided better or at par predictive performance than existing RA methods. The better performance of the SRA method suggests that it may be more reliable than existing RA methods in identifying the target features. Further, unlike the simulated data results, different SRA methods have shown different performances. In the case of Study I and Study II, SRA-Ridge has the best predictive performance, but in Study III, SRA-Lasso has the best predictive performance, which suggests that SRA methods performance may change with dataset and ensemble models. In general, the variation in performance of feature selection techniques with dataset has been reported in the literature and could be attributed to data characteristics [40].

**Table 4:**
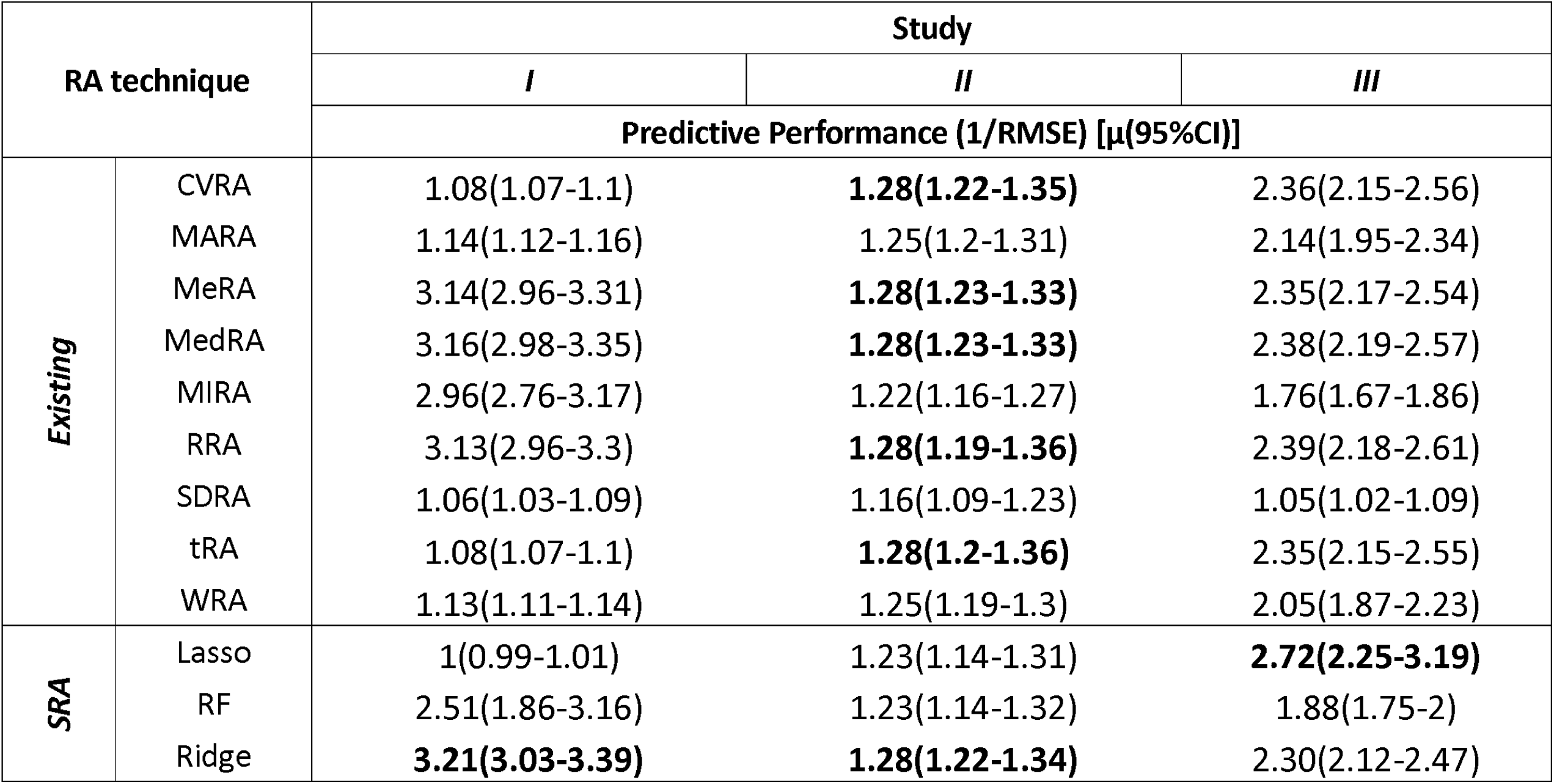
Comparison of SRA methods with Existing methods for three real studies in terms of outcome prediction (1/RMSE)

In the current study, Study III data is also used to compare the performance of SRA based selected methylated features with state-of-art literature based selected features [41,42]. SRA-Lasso is used to obtain the target features. The complete dataset is used for the FS step rather than the training data. SRA-Lasso identified 484 methylation sites compared to 353 methylation sites identified by the literature, but only ten methylation sites are shared between the two approaches (Supplementary File 1). The selected methylation sites from the two approaches are compared for their predictive performance on the test data. Accordingly, the Study III dataset is split into training (80%) and test (20%) data. RIDGE model is prepared using training data followed by predictive performance measurement on test data. It is found that SRA-Lasso based selected features provided a marginally better predictive performance (RMSE^-1^ (95% CI): 0.06 (0.05-0.07)) compared to literature recommended selected features (RMSE^-1^ (95% CI): 0.05 (0.04-0.05)).

Further, we identified the differentially expressed genes associated with selected methylated sites using *BioMethyl* package [43]. SRA-Lasso based selected methylated sites are linked with 288 genes, but literature based selected methylated sites are linked with only 136 genes (Supplementary File 2). Only ten genes, namely SFRP1, STRA6, BNC1, CSPG5, DCHS1, DIRAS3, TCF15, ERG, PIPOX, and MCAM, are shared between the two approaches. Literature also provides a database, GenAge, of 307 genes commonly associated with age [44]. Among the 136 genes linked with literature-based methylation sites, only 1 out of 308 genes is found (Supplementary File 3). However, among the 288 genes linked with SRA-Lasso based methylation sites, 9 out of 308 genes are found (Supplementary File 3). Thus, SRA-Lasso may be relevant in identifying target features that have both biological importance and good predictive performance.

## Conclusion and Discussion

This paper proposes SRA, an innovative approach, to perform rank aggregation in ensemble models for feature selection. The approach allows dynamic learning of feature performance pooling strategy, which current rule-based rank aggregation methods do not perform. The approach is flexible and could be incorporated into any ensemble technique. The SRA could identify target features while retaining very few noise features compared to other methods. The simulated data studies showed that SRA outperforms existing methods in feature selection and prediction performance. Similar performance in real datasets also demonstrates the practical relevance of SRA.

The proposed method has certain limitations. The scope of the current study is limited to concept testing. Consequently, the robustness of the approach on different data types and modeling techniques could be the focus of future research. The ensemble model used in the study assumes a linear combination of features. Thus, future research could study SRA for algorithms designed to explore the non-linear combinations of features.

## Key Points

- Supervised Rank Aggregation (SRA) methods are better than rule-based rank aggregation methods for ensemble-based feature selection (EFS).
- SRA Ridge could give much better discrimination between true and noise features as well as predictive performance than rule-based rank aggregation methods
- SRA could be useful in detecting the genomic features like methylation sites which could have biological relevance

## Declarations

### Ethics approval and consent to participate

Not Applicable

### Consent for publication

Not Applicable

### Availability of data and materials

All the datasets and code are in the github link: https://github.com/rahijaingithub/SRA.

### Author Contributions

ALL AUTHORS HAVE READ AND APPROVED THE MANUSCRIPT.

**Conceptualisation:** RJ, WX

**Formal Analysis:** RJ

**Investigation:** RJ

**Methodology:** RJ, WX

**Software:** RJ

**Supervision:** RJ, WX

**Validation:** RJ, WX

**Writing-original draft**: RJ

**Writing-review & editing**: RJ, WX

## Acknowledgements

Not Applicable

